# Miniaturizing wet scrubbers for aerosolized droplet capture

**DOI:** 10.1101/2021.03.23.436614

**Authors:** Ulri N. Lee, Tammi L. van Neel, Fang Yun Lim, Jian Wei Khor, Jiayang He, Ravi S. Vaddi, Angelo Q.W. Ong, Anthony Tang, Jean Berthier, John S. Meschke, Igor V. Novosselov, Ashleigh B. Theberge, Erwin Berthier

## Abstract

Aerosols dispersed and transmitted through the air (e.g., particulate matter pollution, bioaerosols) are ubiquitous and one of the leading causes of adverse health effects and disease transmission. A variety of sampling methods (e.g., filters, cyclones, impactors) have been developed to assess personal exposures. However, a gap still remains in the accessibility and ease-of-use of these technologies for people without experience or training in collecting airborne samples. Additionally, wet scrubbers (large non-portable industrial systems) utilize liquid sprays to remove aerosols from the air; the goal is to “scrub” (i.e., clean) the exhaust of industrial smokestacks, not collect the aerosols for analysis. Inspired by wet scrubbers, we developed a device fundamentally different from existing portable air samplers by using aerosolized microdroplets to capture aerosols in personal spaces (e.g., homes, offices, schools). Our aerosol-sampling device is the size of a small teapot, can be operated without specialized training, and features a winding flow path in a supersaturated relative humidity environment enabling droplet growth. The integrated open mesofluidic channels shuttle coalesced droplets to a collection chamber for subsequent sample analysis. Here, we present the experimental demonstration of aerosol capture into water droplets. Iterative study optimized the non-linear flow manipulating baffles and enabled an 83% retention of the aerosolized microdroplets in the confined volume of our device. As a proof-of-concept for aerosol capture into a liquid medium, 0.5-3 µm model particles were used to evaluate aerosol capture efficiency. Finally, we demonstrate the device can capture and keep a bioaerosol (bacteriophage MS2) viable for downstream analysis.

## Introduction

We inhale a variety of aerosols (e.g., dust particles, ultrafine particulate matter (PM), biologically derived aerosols, and potentially infectious bioaerosols) on a daily basis.^1-3^ When aerosols come into contact with the incredibly large surface area of the lungs, it can trigger and exacerbate allergic reactions, cause serious infections, and induce severe toxic reactions, particularly in immunocompromised individuals.^3-5^ Despite years of research on the effects of aerosols on human health, our understanding of the extent specific types of aerosols can have on our short- and long-term health has focused largely on a number of salient aerosol types (e.g., cigarette smoke, PM2.5 particles)^6,7^ and environmental settings (hospitals, farms, etc.).^8-11^ This is in part due to the fluctuating and highly personalized nature of an individual’s exposure as the formation, dispersion, and transport of aerosols is influenced by both physical (e.g., size, shape, density of aerosols) and environmental (e.g., air currents, humidity, temperature) factors.^12,13^ Coupled with the limitations in personal sampling and ability to analyze chemical and biological composition of the collected sample, researchers and clinicians are still unable to pinpoint the patient-specific triggers for the millions of people worldwide affected by environmentally-induced respiratory illnesses. There is a need for a portable and easy-to-operate air-sampling device to inform individuals of their environmental exposures and provide information on personalized health triggers (Figure 1). In this work, we developed a lightweight and low-cost portable device that can be used without technical training; it utilizes liquid to capture aerosols for subsequent analysis—effectively miniaturizing technology used in industrial wet scrubbers. We aimed to maximize the time aerosolized microdroplets interact with aerosols to improve capture.

**Figure 1.**
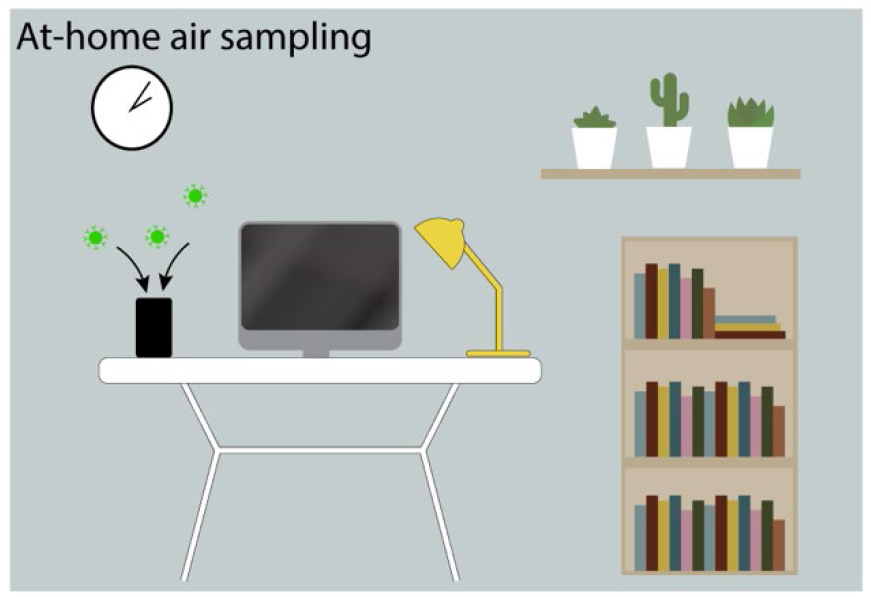
Schematic representation of our battery-powered air-sampling device (small black box to the left of the computer) collecting aerosols in a home environment to provide information on the landscape of aerosols in the room. The concept is that our device can be placed in any environment (e.g., homes, schools, hospitals, playgrounds, farms) and then sent back to a lab for analysis. See Figure 2 for a photograph of the battery-operated, compact device.

To date, a variety of passive^14^ and active^15,16^ sampling devices have been developed for the collection of bioaerosols, including filter collection, centrifugal collection, electrostatic precipitation, liquid impingement, and impaction and direct collection onto growth media.^17-21^ Information on aerosol concentration and size distribution can also be obtained in real-time using aerosol particle counters,^22-24^ however, these monitors do not collect particles nor retain viable samples for further specific identification and analysis.^25^ With respect to direct personal exposure monitoring, some of the aerosol collection methods may not be practical; for example, filter collection and analysis is limited by high elution volumes, bulky setup, and high-power requirements. Due to the development in miniaturization of sampling pumps and electronics, exposure measurement methods including biological aerosol collectors^26-30^ have rapidly advanced and gained popularity.^16,22,31-33^ For non-spore forming infectious bioaerosols (e.g., viruses and vegetative bacterial cells), desiccation during sampling is problematic for downstream viability analyses and infectivity studies.^34^ Although bulk liquid and hydrated gel components integrated in established sampling devices have been shown to improve viability and culturability,^35^ their large collection volume and substantial power requirements limit their applicability in personal samplers and analysis techniques where small collection volume and direct integration with microfluidic devices is desired.

An alternative to collection into bulk liquids are liquid droplet sprays similar to ones used in wet scrubbers.^36^ These systems have been most effective for removing aerosols larger than 1 μm from the air, but they are large (ranging from the size of a small appliance to an industrial smokestack) and have high energy consumption.^37^ The physical phenomena governing the capture of aerosol by droplets in gas flows has been simulated for wet scrubbers^38-40^ and has shown that it is possible to increase the capture efficiency of aerosols (including those smaller than 1.0 µm) by decreasing the droplet size and increasing droplet residence time in the scubber.^40^ The overall collection efficiency can be approximated by summing the modes of capture—diffusion, interception, inertial impaction, and gravitational settling^39^—though most of these require large distances of timescales to operate effectively, which is difficult to achieve in a small scale device. The relationship between aerosol size and predominant mode of capture has previously been evaluated; impaction was found to be a predominant mode for the capture of aerosol sizes (d_p_) > 5 µm, and diffusion for d_p_ < 1 µm.^38^ At d_p_ < 0.05 µm, impaction and interception are negligible and diffusion can be considered as the sole mode of capture. Interception is unaffected by flow rate, but changes with droplet size and packing.^41^ Industrial wet scrubbers generate the required liquid droplets using large spray towers. Alternatively, an attractive method for generating liquid microdroplets is ultrasonic atomizers due to their compact size and use in existing consumer products such as industry for home humidifiers. Recent studies using ultrasonic atomizers to generate sprays of liquid microdroplets showed a decrease in PM2.5 and PM10 concentration over time when the microdroplets were injected near burning incense.^42^

In this work, we harnessed the benefits of microdroplet liquid sprays to fill the need for an accessible and portable, at-home air-sampling device that collects aerosols in a liquid compatible with downstream analysis methods. To our knowledge, our device is the first battery-powered air-sampling device that uses aerosolized microdroplets to capture aerosols for analysis. We demonstrate the function and performance of the device using model fluorescent aerosol. Our results inform how the approach can be used for bioaerosol collection. The portable air-sampling device generates a mist of 4 μm liquid droplets and mixes it with aerosols. The device guides the mixture in a non-linear path to keep microdroplets suspended in the open spaces of the device for as long as possible. The droplet residence time (time the droplet spends suspended in a gas flow) along with droplet packing (number of droplets present) has been shown to increase capture efficiency.^40^ We demonstrate that aerodynamic features enable the capture of the generated mist in a reduced volume and present optimization criteria for device geometry. By leveraging the principles of open microfluidics, we created a pathway via horizontal and vertical open mesofluidic channels for the coalesced droplets to travel and collect for future downstream analysis.^43,44^ We evaluate the effect of flow patterns on droplet retention efficiency and the model aerosol capture efficiency for a 0.5-3.0 µm size range. Finally, we demonstrate the device can capture and keep a bioaerosol viable for downstream analysis.

## Experimental section

### Device fabrication

The body of the device and ultrasonic atomizer cup were designed in Solidworks 2017 and 3D printed out of black resin (RS-F2-GPBK-04) using a Form 2 or Form 3 stereolithography 3D printer (Formlabs). The parts were cleaned in isopropyl alcohol (IPA) for 10 minutes in a FormWash (Formlabs) and rinsed down with clean IPA to remove excess uncured resin. The devices were dried with compressed air and cured under a 395-405 nm 20W UV lamp (Quans) for 1 hr. A 1.3 mm PS thick window was milled (Datron Neo) to enable visual access to the capture region of the device.

### Portable Electronics (Figures 2A and 6)

The ultrasonic atomizer (Comidox) with a frequency of 113 KHz and 730 apertures 5 µm in diameter was adhered to the floor of the ultrasonic atomizer cup with 100% silicone caulk (Gorilla Glue). A 40 mm square computer fan (Model OD4010-05HB55, Orion Fans) was used to generate airflow. The voltage delivered to the fan was controlled by a microcontroller (Arduino Micro, Arduino). A darlington transistor (TIP122, STMicroelectronics), 100 Ohm resistor (SparkFun Electronics), 22 and 24 AWG jumper wires (OSEPP Electronics LTD), and 2.1 mm male and female barrel jacks (Centropower) were used. The device was powered by a 9V battery (Zeus Battery Products).

### Simulations (Figure 5)

Computational Fluid Dynamics Module in COMSOL Multiphysics v5.5 software was used to simulate the air flow and pressure in different device geometries in a stationary study. A 2D cross-section of our 3D device was modeled. The *Laminar Flow* physics interface, governed by the Navier-Stokes equations, was selected to model air flow in the device based on the estimated Reynolds number calculated (Re<1200). Two fan inlets with a fully developed flow profile, initial flow rate = 6.3 L/min, and entrance thickness of 0.02 m was used. An outlet boundary was defined by P=0 Pa with backflow suppressed. No slip boundary conditions were applied to all remaining boundaries. A fine element quadrilateral mesh with extra fine element boundary refinement at the baffles was used.

### Droplet retention gravimetric analysis using non-portable electronics setup (Figure 6)

Droplet retention was measured by weighing the ultrasonic atomizer cup filled with water and the 3D printed body separately using an analytical balance (Mettler Toledo ME103E) before and after operating the device for 30 seconds with the fan operating at 25% voltage for an airflow rate of 6.3 L/min.

### Aerosol capture efficiency (Figure 7)

A testing chamber (0.56 × 0.52 × 0.42 m) was used to determine the capture efficiency of our device using fluorescent polystyrene latex (PSL) particles (Fluoresbrite® YG Microspheres, Polysciences Inc.); particle sizes tested: 0.5, 0.75, 1, 2, and 3 μm. The particles were diluted in DI water and aerosolized using a VixOne Nebulizer (Westmed Inc.); each particle size had a dedicated nebulizer to prevent contamination of other particles sizes. An aerodynamic particle sizer (APS 3321, TSI Inc.) was used to monitor the particle size aerosolized and particle concentration during experimental runs. Particle concentration varied by size; see Table S1 for more information. Three open-face aerosol reference filter holders (EMD Millipore, Model #XX5004710) with 0.2 μm PTFE Omnipore membrane filters (Millipore Sigma, product #JGWP04700) were co-located with our air-sampling devices in the chamber to collect particles; filter membranes were collected for analysis Flow rates for reference filters placed in the chamber were calibrated using a mass flow meter (TSI Inc., Model #4140) to 3.15 slpm (half the flow rate of our device due to limitations in the flow controller). A humidity monitor [(Extech Instruments, Model #SD700; AcuRite, Model #01083M)] and a dry airline in the chamber was used to keep the humidity between 60-70%. An interchangeable particle capture region was pre-wetted before placing the fully assembled devices into the chamber; the sampling region in the device was replaced between chamber runs. Devices and reference filters captured particles for 25 mins followed by a 3 min chamber purge before devices being removed from the chamber. A 1 oz. polystyrene cup collected the sample (Uline, product #S-14487) and was weighed (Smart Weigh Pro Pocket Scale TOP500, Amazon) before and after chamber runs. Sampling regions were rinsed down with approximately 5 mL of DI water after chamber runs. All samples were diluted to 25 g (25 mL) using DI water which matched the volume of water used for the reference filters. Reference filters were carefully placed in 50 mL polypropylene centrifuge tubes (ThermoFisher, product #339652) with 25 mL DI water.

### Fluorescence measurements (Figure 7)

Capture efficiency was calculated as 100 x (fluorescence of samples from our device/fluorescence of co-located reference filter). Fluorescence measurements were performed using a fluorometer set to a gain of 200 (Sequoia-Turner model 450). Sample cups and centrifuge tubes containing the reference filters were vortexed (Vortex-Genie 2, Scientific Industries, Inc.) for at least 8 mins at 1800 rpm and remained vortexing until analysis. A 5 mL aliquot of the sample was transferred to a 12×75 mm borosilicate disposable culture tube (Fisherbrand, Cat No. 14-961-26) and discarded to rinse the tube of any previous sample. The sample was re-aliquoted into the test tube and measured then discarded, for a total of three measurements. Between each aliquot, the pipette was used to resuspend the particles in the sample cup once before measurement. A new glass test tube was used for each particle size on any given day. Blank measurements were always taken when switching to a new glass test tube. Calibration curves for each particle size are available in Figure S5. Blank measurements were subtracted from sample and reference fluorescent values prior to additional calculations and statistical analyses.

### MS2 aerosolization and analysis (Figure 8)

A testing chamber operating similarly to the one described previously was used to determine if our device was capable of capturing a bioaerosol (bacteriophage MS2 ATCC #15597-B1; see SI for more information on propagation). A 1 mL MS2 solution (1.9×10^11^ PFU) was aerosolized (VixOne Nebulizer) in a closed chamber. An aerodynamic particle sizer (APS 3321, TSI Inc.) was used to monitor the particle concentration during experimental runs. A 25 min sampling period followed by a 5 min purge was used. Device samples collected in the 1 oz. polystyrene cups were collected with no further dilution. A double agar layer plaque assay was performed to assess MS2 viability in a bacterial host (*Escherichia coli*). Briefly, Tryptic Soy Agar (TSA) (BD Difco #236950) was prepared following manufacturer’s directions, added to petri dishes (Fisherbrand 100×15mm #FB0875712), and allowed to cool at room temperature. A ten-fold serial dilution was performed using 1X PBS for all samples. 100 µL of diluted sample and 100 µL of an *E. coli* Famp (ATCC #700891) suspension were added to 7 mL of top agar [0.5% (w/v) NaCl (Fisher Scientific #S271-500) and 0.7% (w/v) Bacto Agar (BD #214010)] in a borosilicate glass tube (Fisherbrand #14-961-27) after which the solution was mixed by rolling tube between hands and subsequently poured onto prepared TSA plates; top agar was cooled before plates were inverted and incubated overnight at 37 °C. Plaques were counted the following day. The assay was performed in duplicate for all sample dilutions, including undiluted samples. Additionally, a negative control (only *E. coli*) and PBS control (PBS added in place of sample) were also included.

### Data analysis

All data and statistical analyses were performed using Prism v9.0 (GraphPad) software.

## Results and Discussion

### Design considerations

We developed a portable air-sampling device, approximately the size of a small teapot, that can be placed in many environments (homes, schools, hospitals, playgrounds, farms) for aerosol capture. Major considerations for the device included how to keep the device within a comfortable size to place on a desk, battery-powered to avoid the need of a power outlet, and simple to operate in a variety of environments. An additional consideration included maintaining the captured bioaerosols in liquid to ensure their viability and compatibility with rapid downstream analysis, a point of interest for future studies. While we focused the demonstration herein on manipulating airflow for microdroplet retention, we also used model inert particles to demonstrate the capability of the air-sampling device with bioaerosol applications in mind. An air-sampling device based on aerosolized microdroplets has the potential to capture a wide range of aerosols^37^ and is amenable for use with a variety of liquid capture solutions (i.e., water, surfactant, media, and solvents). Given the efficacy of wet scrubbers, we chose to incorporate liquid droplets in our portable sampling device. Our air-sampling device consists of a small fan, an ultrasonic atomizer (droplet generator), and a 3D printed body (Figure 2A). The fan draws in air from the environment, and an ultrasonic atomizer generates 4 μm liquid droplets which move in the same direction as the air flow; the goal is for the liquid droplets to capture aerosols that enter throughthe fan. In our device, the microdroplets have an initial velocity that is independent of the air flow generated by the fan. Although the fan flow rate is tunable by adjusting the voltage delivered to the fan, the aerosol experiments and simulations in this manuscript were conducted with a flowrate of 6.3 slpm. Additionally, the initial velocity of the microdroplets is constant due to the designed frequency of the ultrasonic atomizer. The microdroplets and aerosols are carried in a non-linear path by the airflow generated by the fan and coalesce on fluidic baffles which will be discussed further in Figures 3 and 4. The baffles contain open mesofluidic channels that guide the coalesced droplets to a collection reservoir at the bottom of the device (Figure 2B). The clean liquid reservoir and sample collection reservoir were designed to each hold up to 200 mL of fluid for an operation time of 1 hour but the device is amenable to other volumes and operation times (Figure 2A). Electronic components (fan and ultrasonic atomizer) were made compatible for battery operation to improve overall portability, but the battery also limits the operation time (Figure S1). Due to the low pressure drop of the flow path, the device can be operated using low-cost fans, i.e., does not require pumps, typical for most of the widely used air-sampling devices today (e.g., Anderson cascade impactors, Burkard personal volumetric air sampler, Coriolis biological air sample, cyclones, and impingers).

**Figure 2.**
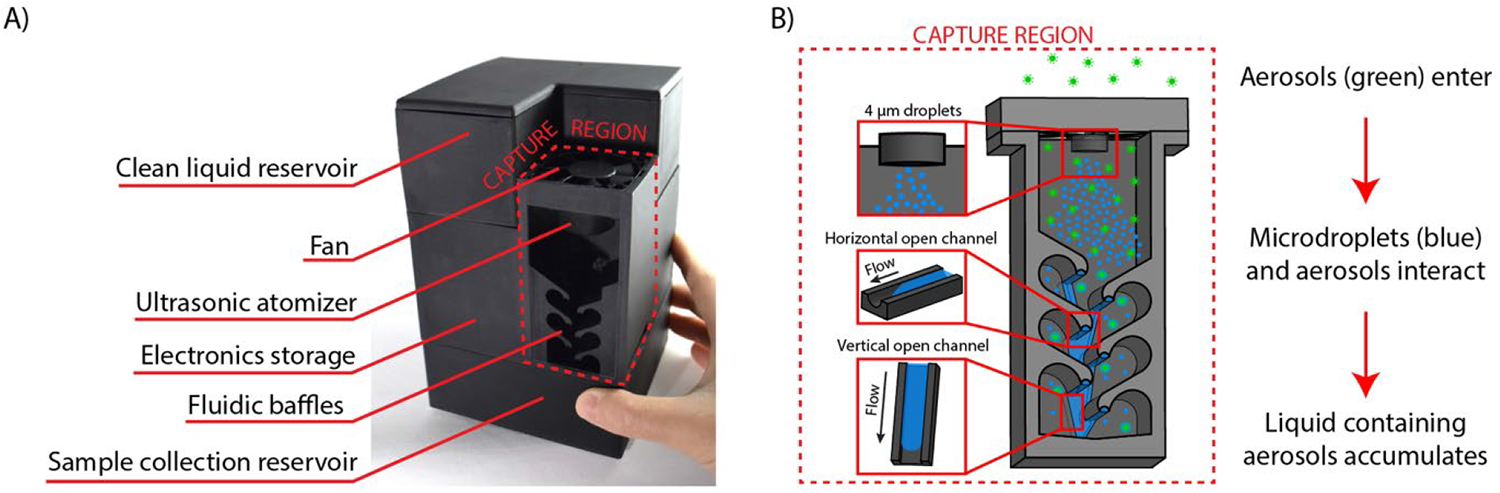
Portable droplet-based air-sampling device. (A) Photograph of device. (B) Schematic of capture region workflow. An ultrasonic atomizer generates 4 µm droplets (top), and horizontal (middle) and vertical (bottom) open mesofluidic channels collect coalesced droplets and aerosols. Aerosols enter through the fan, microdroplets intercept the aerosols, and six angled baffles guide the airflow to increase capture of aerosols. Front wall of the device has been removed for visualization of the device.

**Figure 3.**
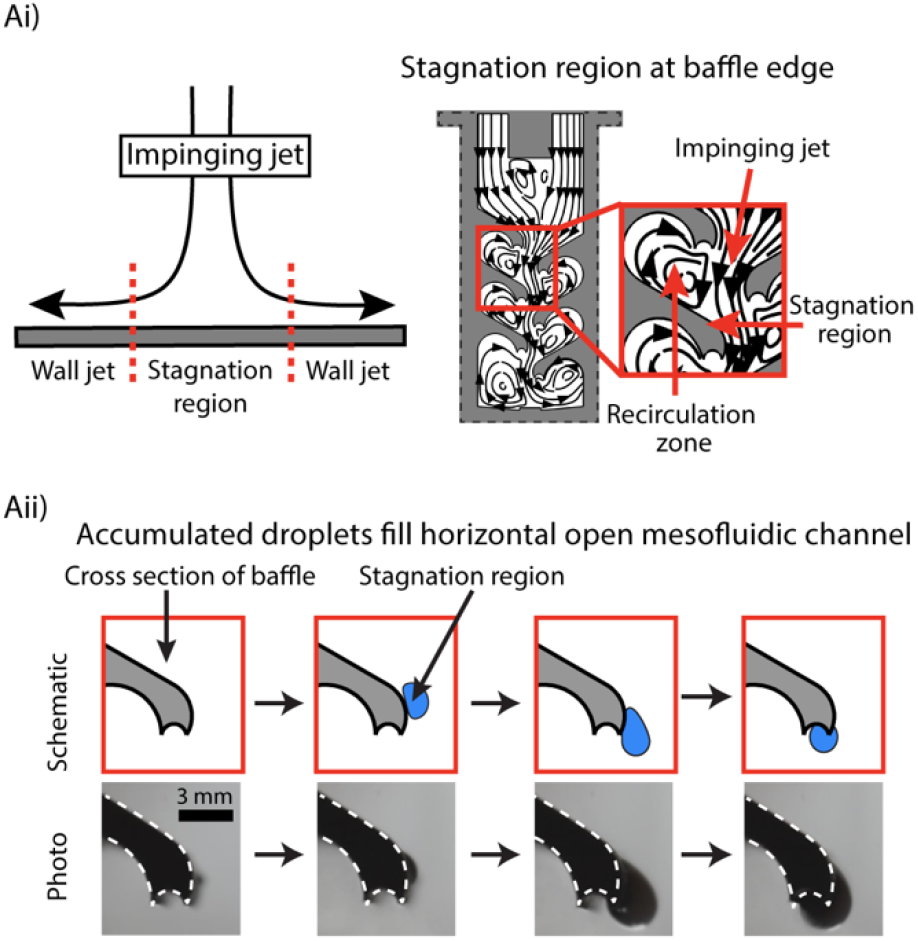
Impinging jet, stagnation region, and horizontal mesofluidic channels. (Ai) Basic schematic representation of an impinging jet (left) and the stagnation region in the device (right). (Aii) Droplets accumulate at a stagnation region on the baffle and migrate down the edge of the baffle, where it travels into the horizontal and vertical channels. Images were taken with a device that did not have a back wall, and water was colored dark blue to enable better visualization of the droplet. Video of droplet formation at the stagnation region is available in Supplementary Video 1.

**Figure 4.**
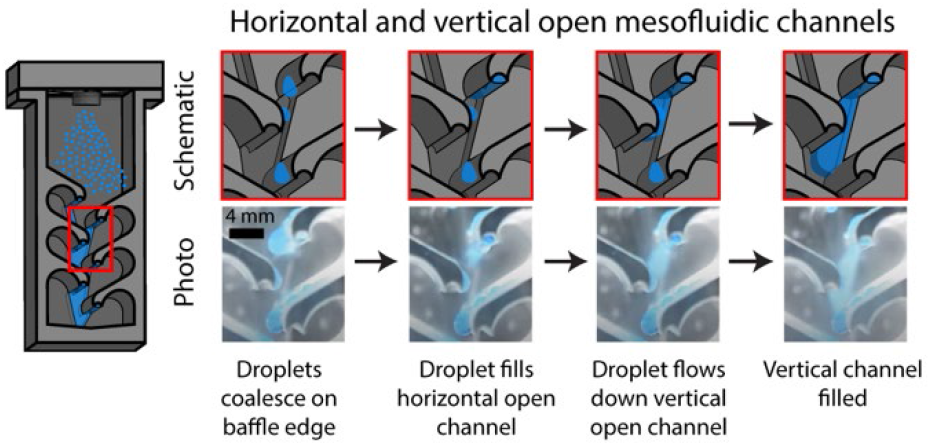
Schematic (top) and photo (bottom) progression of filling the mesofluidic channel at the edge of a baffle. Video was taken using a clear device to enable better visualization of the channel and is available in Supplementary Video 2.

### Flow guiding baffles and open fluidic channels

The capture region of the air-sampling device was designed to increase interactions and mixing of microdroplets and aerosols. In traditional wet scrubbers, the droplet paths are linear, so to increase microdroplet residence times, it is necessary to increase the height of the device. In order to minimize the device size and maintain portability, we designed fluidic baffles that generate recirculation zones with longer residence time than the main airflow (Figure 3Ai). Air and droplets follow the flow paths and populate regions under each baffle due to a low Stokes Number (<0.03). Internal geometry of the flow channel consists of a series of horizontal open channels underneath the baffles, connected by a vertical open channel along the back wall (Figure 2B). The constriction bends to form an impinging jet on the upper surface of each baffle (Figure 3). The behavior of impinging jets has been described in the literature^45-47^ and their comprehensive analysis is beyond the scope of this paper. However, for the purpose of this work, it is important to understand that angled impinging jets create stagnation regions (Figure 3Ai, left).

In our device, the stagnation regions occur at the edge of the baffles; a portion of the jet follows the curve of the baffle (creating recirculation zones), and the remaining jet joins the bulk flow in the constriction (Figure 3Ai, right). Microdroplet growth and transition to liquid effluent is likely due to two complementary mechanisms: (i) microdroplet coalescence and (ii) heterogeneous growth in a supersaturated environment. These are similar to the growth of combustion-generated particles, where recirculating leads to the formation of large super-aggregates.^48,49^ It is challenging to model these processes, however, the highest probability of droplet coalescence and heterogeneous growth occurs in regions with long residence time and high droplet concentration. In our geometry, these conditions are present at the stagnation regions and in the recirculation zone. Once larger droplets are formed, they either settle on the surface due to gravity or via inertial impaction associated with the impinging jet. On the surface, these larger droplets migrate to the edge under the aerodynamic load acting on the droplet^50-53^ or by gravity and are collected in the horizontal open mesofluidic channel embedded in the edge of the baffle (Figure 3Aii). The horizontal open channel guides the aerosol-laden sample toward the back wall where it meets the vertical open channel; this vertical open channel connects all of the baffles and drains the sample into a collection reservoir (Figure 4).

### Computational modeling of airflow and effects of baffle geometry

Computational modeling aimed to (1) better visualize formation of recirculation zones and air flow around the baffles and (2) improve droplet retention within the device (reducing droplet loss through the outlet). The microdroplet-aerosol collisions were increased by incorporating baffles thus promoting the formation of recirculation zones. The multiple impinging jet flow pattern enables inertial impaction of droplets onto surfaces for capture. Using Computational Fluid Dynamics (CFD) COMSOL Multiphysics), we performed a parameter space optimization of the features in the capture region, specifically looking at how they affected airflow (Figure 5), and ultimately improved droplet retention (Figure 6). We solved for a mean air velocity field and pressure at a steady-state. We modeled a 2D cross-section to reduce computational load.^54^ Recognizing the challenges associated with modeling and validation of aerosol-laden flow,^55^ here CFD was used as a comparative tool to inform device design iterations and not as an absolute or quantitative method. The full details of the model are presented in the experimental section. Briefly, the following boundary conditions were used: a fully developed flow rate profile was used for the inlet with a zero-pressure outlet; all other walls were modeled with a no-slip boundary condition.

**Figure 5.**
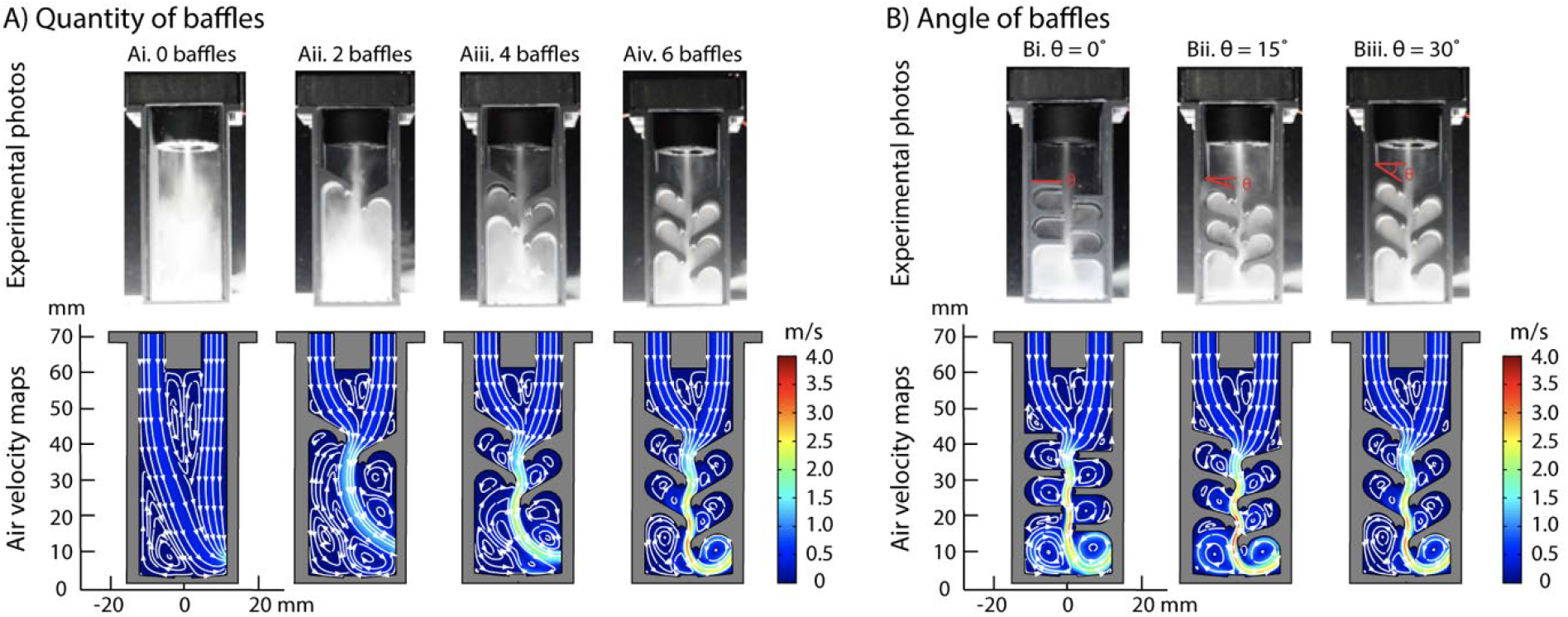
Effect of baffle quantity and angle on airflow. Experimental photos showing droplets in device and mean airflow velocity maps of a 2-D cross section of the device with (A) 0, 2, 4, and 6 baffles (all at a 30° angle) and (B) 0°, 15°, and 30° baffle angles (Supplementary Video 3). Images were taken from different timepoints and should not be compared with one another. Colors represent velocity and indicate the fluid flow (airflow) modeled. White lines and arrows represent direction of flow. Devices used in photos do not have front or back walls to enable visualization of the droplets.

Experimentally, we observed droplets accumulating on all surfaces in the device, coalescing to form larger droplets, and being shuttled away by the mesofluidic channels to the collection area (Supplementary Video 1 and 2). Modeling the mean airflow in the device enabled us to extract details such as the direction and pattern of the airflow; the flow within the device is laminar with a calculated Reynolds Number of less than 1200. To understand the effect of baffle geometry on recirculation zone formation and air flow, we varied the quantity and angle of the baffles (Figure 5). More recirculation zones were created with each additional set of baffles (Figure 5A). The baffles also increased the probability of droplet capture due to (i) increased surface to volume ratio and (ii) greater number of impinging jet regions where the high velocity interacts with the wall. This is supported by experimental droplet retention results shown in Figure 6. Further, each extra set of baffles constricts the main airflow resulting in an increased velocity of the impinging jet, allowing for more effective liquid transport from the stagnation regions to the mesofluidic channel (Figure 3). The horizontal space between the baffles controls the airflow constriction, but if too small can also cause backflow in the device. The number of stagnation regions increases with the quantity of baffles, providing more areas of high coalescence for droplets (Figure 3A). Finally, the pressure above the top baffles increases with each additional set of baffles and decreases as the angle increases (Figure S2). Based on these observations, the six-baffle design was used in aerosol capture efficiency studies to be discussed in Figure 7.

### Microdroplet retention efficiency

The microdroplets generated to capture aerosols in air must subsequently be captured by the device to perform analyses. The microdroplet capture process is challenging due to the high rate of air flow through the device and the need to not constrict that air flow. As the ultrasonic atomizer generates microdroplets and the fan draws aerosols into the device, an airflow exhaust must be designed to prevent flow exiting through the inlet in the case of high back pressure. An unobstructed geometry will lead to microdroplets flowing through the device without being captured. We sought to maximize the microdroplet retention while keeping an outlet to prevent backflow. The pressure drop associated with the impinging jet at stagnation regions helps coalesce the droplets on the baffles, however, it should not be high enough to cause backflow. We chose not to employ a mesh filter to increase microdroplet retention because recovering bioaerosols from filters can be damaging.^56^ Additionally, droplets clog the small apertures in mesh surfaces which blocks airflow. Through various design iterations, we measured the percentage of liquid retained by weighing the ultrasonic atomizer filled with water and the aerosol capture region of the device before and after operating the ultrasonic atomizer. Microdroplet retention increased with numbers of baffles with an 81% droplet retention in the six-baffle design (Figure 6Aiv). We also studied the effect of the angle of the baffle on microdroplet retention. There was an increase in microdroplet retention between 0° and 15° baffle angle followed by a decrease between 15° and 30° baffle angle. A 15° baffle angle had the largest microdroplet retention of 83%, while 0° and 30° baffle angle had a 73% and 81% microdroplet retention, respectively (Figure 6B). Since there was not a statistically significant difference in microdroplet retention between a 15° and 30° baffle angle, we chose to move forward with the 30° baffle angle due to the reduced pressure near the fan inlet, which is favorable in reducing backflow as discussed previously (Figure S2).

**Figure 6.**
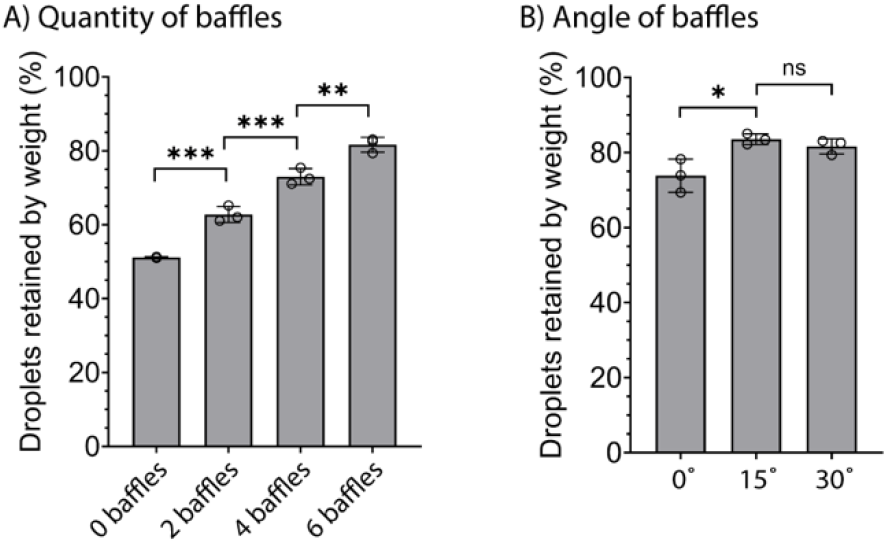
Effects of baffle quantity and angle on microdroplet retention. Microdroplet retention trends were determined by weighing liquid retained in the device before and after running the ultrasonic atomizer (see Experimental section). (A) The six-baffle design has the highest microdroplet retention (81.6% ± 1.6%) and was used for further device iteration studies. (B) The 15° baffle angle design retained a higher percentage of microdroplets (83.5% ± 1.1%) than a 0° baffle angle (73.8% ± 3.6%). No significant difference (ns) was observed with an additional 15° baffle angle (30°; 81.6% ± 1.6%). Bar graph represents the mean ± SD of n=3 experimental tests. One-way ANOVA with post-hoc Tukey’s multiple comparison test; * p<0.05; ** p<0.005; *** p<0.001. Devices tested in (A) all had a 30° baffle angle.

### Aerosol capture efficiency

An aerosol chamber^57^ described previously was used to test the six-baffle, 30° baffle angle portable device for aerosol capture efficiency. Briefly, monodispersed fluorescent polystyrene latex spheres of varying sizes (0.5, 0.75, 1, 2, and 3 μm) were aerosolized in separate experiments using a nebulizer (Table S1). Two fans placed in opposite corners of the chamber were used to provide well-mixed conditions in the chamber. Three devices, co-located with reference filters, were tested in the chamber at the same time for 25-minutes. The reference filter flow rate matched the measured flow rate of our device (Figure S3). 25 mL of DI water was used to elute the particles from the filter and additional DI water was added to the device sample until the volume was also 25 mL. We measured the fluorescence of the liquid sample and used a ratio to determine aerosol collection efficiency. Particle sizes were selected based on model particles commonly used in bioaerosol research. The use of model polystyrene latex spheres for device testing was to avoid unnecessary exposure to bioaerosols (Table S2). It is important to note that we did not expect to reach 100% capture efficiency, nor is this required for analysis; further for bioaerosol detection, amplification steps (such as culture and PCR) can be employed. The highest efficiency observed in our device was 17.5% with 0.5 μm particles and the lowest efficiency was 4.5% with 2.0 um particles (Figure 7). Differences in aerosol capture efficiency were observed between the three devices tested, likely due to heterogeneity in the microdroplets produced by the ultrasonic mist generators (Figure S4) and inherent differences in the 3D printing of the devices. The reproducibility across devices could be improved with more accurate fabrication methods. Importantly, we note that variability in capture efficiency was also observed in reference filters used likely due to the manual control of the flow rate and particle nebulization (Table S3). Based on prior literature for other capture devices,^58-61^ aerosol capture efficiency often varies with each particle size tested.

**Figure 7.**
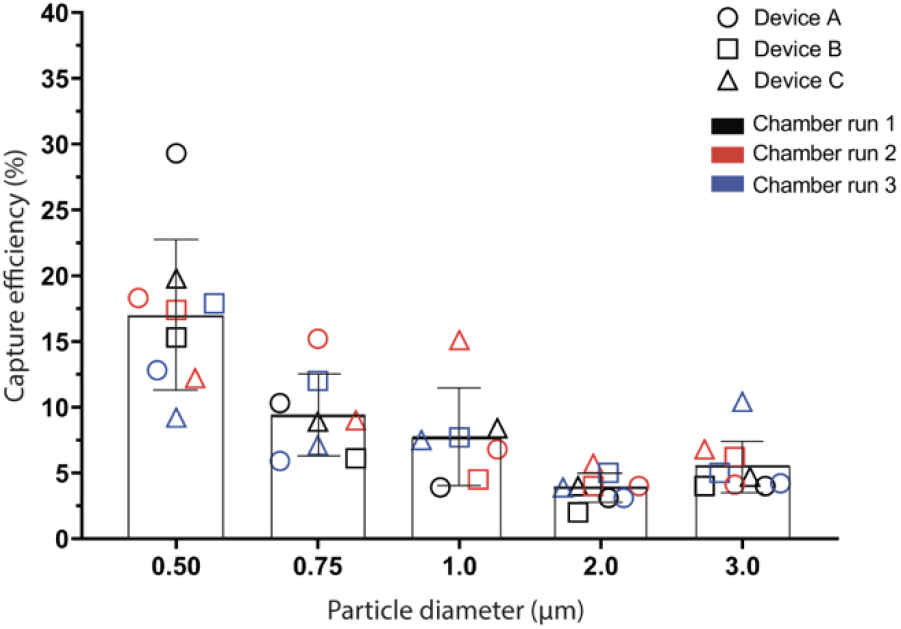
Capture efficiency of our portable air-sampling device (n=3 devices) for 0.5, 0.75, 1.0, 2.0, and 3.0 μm polystyrene particles. The highest aerosol capture efficiency observed was 17.5% with 0.5 μm particles and the lowest capture efficiency observed was 4.5% with 2.0 μm particles. Bar graphs represent mean ± SD of n=3 independent chamber runs for each device. Symbol shape indicates different device A, B, or C; symbol color indicates independent chamber runs.

### Bioaerosol capture

To demonstrate a potential application of our device in capturing bioaerosols while maintaining viability, we aerosolized a bacteriophage MS2 solution in a chamber with our device and analyzed the collected liquid from the device. MS2 is a virus that infects *E. coli* and is commonly used as a surrogate viral particle in aerosol studies for safety concerns.^62^ Briefly, a MS2 solution was nebulized in a closed aerosol chamber containing a device for a 25-minute sampling period. As a control, we also performed a chamber run where no MS2 was nebulized. We then performed a plaque assay with the liquid samples collected from our device to determine if MS2 was present and viable. Plaque assays are used to quantify infectious viral particles.^63^ The results from the control showed no MS2 (Figure 8, left). In comparison, samples from the chamber runs where MS2 was nebulized show MS2 was captured by the device and able to infect *E. coli*, an indication that the MS2 remained viable (Figure 8, right).

**Figure 8.**
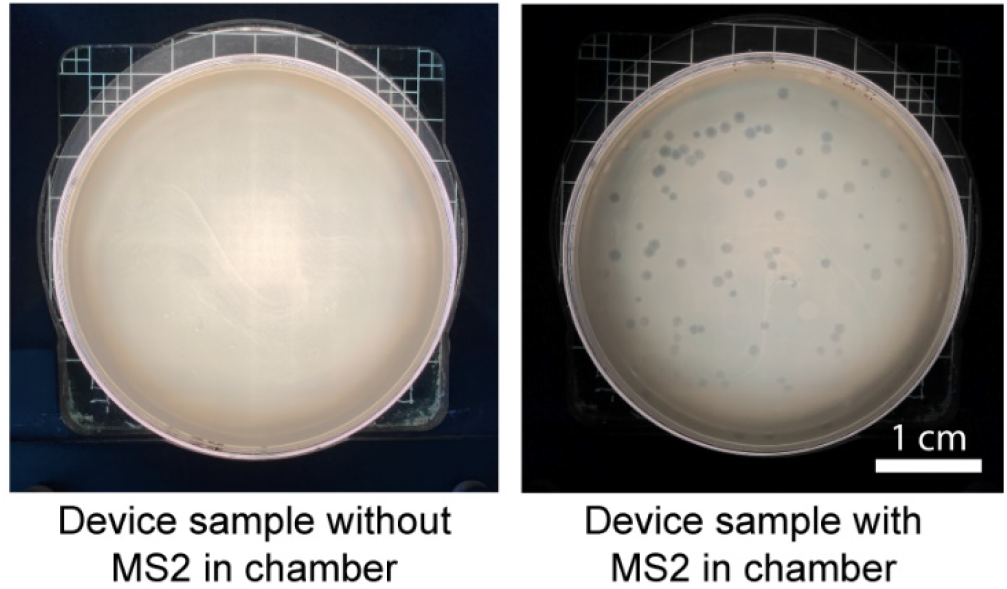
Bacteriophage MS2 captured in our air-sampling device remains viable. A plaque assay was performed using the liquid samples obtained from our device. A control liquid sample obtained from our device (no MS2 aerosolized in chamber) showed no plaques (left) when cultured with *E. coli*, demonstrating MS2 absence from the sample. In contrast, when MS2 was aerosolized in the chamber, plaques formed (dark grey circles; right) as a result of MS2 infecting *E. coli*, indicating MS2 presence in the sample. Representative images of n=2 independent chamber runs with one device.

To our knowledge, our device is the first battery-powered air-sampling device that uses microdroplets to capture aerosols for analysis. The high humidity environment generated within the device coupled with storage of the captured aerosols in a fluid through the duration of the sampling period has the potential to keep bioaerosols viable for downstream culture and analysis; in this case, high capture efficiency is not critical to the identification of the captured bioaerosol because the sample can be amplified via culture (or detection methods like PCR). Additionally, the results were obtained with off-the-shelf low-cost components that can be further optimized to achieve higher efficiencies, however the device in its current state can be used in real-world environments. The bulk fluid also enables seamless integration with existing downstream analysis methods. To collect a sample from their environment, the only steps required by the user are: fill the reservoir with an appropriate fluid, flip the power switch, and put a cap on the collection reservoir after sampling. They would then mail the sample to a lab for analysis. Future work includes developing the sample preservation and analysis pipeline for bioaerosols of interest. We note that there are numerous considerations and validation steps required in future work including selection of media; considerations will vary based on the analysis goal (e.g., simple detection of the presence of a microbe (yes/no) versus quantification). The platform introduced in this manuscript opens many avenues for future investigation.

## Conclusion

In this work, we presented a portable air-sampling device that utilizes aerosolized microdroplets for aerosol capture. Future studies will focus on capture of bioaerosols for downstream analysis, viability studies, and infectivity assays. Our device is also well-positioned to be compatible with other low-viscosity fluids due to the working principles of the ultrasonic atomizer. Maximizing the capture of the liquid droplets after aerosolization by the ultrasonic atomizer will be part of future investigations to ensure the safety of expanding the use of potentially harmful fluids, such as organic solvents, as the capturing fluid. Ultimately our vision is that our air sampling platform will enable people who typically do not have access to or cannot afford services or products that measure environmental exposures (e.g., those in lower socioeconomic groups or high-risk environments), to collect air samples and determine the levels of environmental exposure within their daily lives.

## Supporting Information

The supporting information includes extended materials and methods on the simulations, fan flow rate measurements, electronics set up, and phage MS2 propagation (Figures 3-8). Additional figures, videos of droplet generation, and tables of particle concentrations are also included.

## Acknowledgments

We thank Bret Nestor for helpful discussion designing the electronic components and writing the Arduino code. This work was supported by the David and Lucile Packard Foundation (ABT, UNL, FYL, EB), the University of Washington, National Institutes of Health (NIH1R35GM128648, TLVN; NIHR21ES024715, JH, RSV, IVN) and the Society for Laboratory Automation and Screening (SLASFG2020, UNL). Any opinions, findings, and conclusions or recommendations expressed in this material are those of the author(s) and do not necessarily reflect those of the Society for Laboratory Automation and Screening or the NIH.

## Conflicts of Interest

ABT has ownership in Stacks to the Future, LLC and EB has ownership in Stacks to the Future, LLC, Tasso, Inc., and Salus Discovery, LLC. However, this research is not related to these companies.

