## Supplementary material for "Miniaturizing wet scrubbers for aerosolized droplet capture": SI

Extended Materials and Methods

Contents

Figure S1- Photo of non-portable and portable electronic components
Figure S2- Pressure maps of devices in Figure 5
Figure S3- Measured air flow velocity in non-portable and portable devices
Figure S4- Droplet size distribution of each device used in Figure 7
Figure S5- Calibration curves for each fluorescent particle size in Figure 7

Table S1- Particle concentration in chamber for each particle size
Table S2- Bioaerosol size range
Table S3- Relative standard deviation for reference filters and devices

Videos

V1-Stagnation point and droplet flowing in horizontal channel from Figure 3
V2-Droplet flowing down vertical channel from Figure 4B
V3-Droplet flow in device iterations from Figure 5

### Extended Materials and Methods

**Non-portable Electronics (Figures 3-6):** The ultrasonic atomizer (Comidox) with a frequency of 113 KHz and 730 apertures 5 µm in diameter was adhered to the floor of the ultrasonic atomizer cup with super glue (Ultra Gel Control, Loctite) or 100% silicone caulk (Gorilla Glue). A 40 mm square computer fan (Model AB4010M12**,** HK Fan) was used to generate airflow. The voltage delivered to the fan was controlled by a microcontroller (Arduino Uno, Arduino) and L298N transistor (Qunqi). Two power supplies were required for this setup. Add M/F barrel jacks

**Simulations (Figure 5):** The *Laminar Flow,* physics interface, governed by the Navier-Stokes equations, was selected to model air flow in the device. Air was modeled as an incompressible fluid with standard fluid values at 1 atm (reference pressure) and 293.15 K (reference temperature). Two fan inlets and one outlet were set with the remaining boundary conditions set no slip. A fine element, free quadrilateral mesh was used for the majority of the device; an extra fine, free quadrilateral mesh calibrated for fluid dynamics was used for the baffle features. A stationary study was used with a direct, fully coupled linear solver.

**Fan flow rate measurement (Figure 7):** The device was attached to a 1 ft long pipe and a hot-wire anemometer (AN-1005) is used to measure the velocity profile at the outlet of the pipe. These velocity measurements are also used for calculations of the flow sampling rate. TSI 1213-20 hot wire probe connected to an anemometer is positioned at the outlet. The anemometer is calibrated for the range of 0.2 m/s - 7 m/s using the standard calibration procedure. The data from the anemometer is collected at a frequency of 10 kHz with a data acquisition module (National Instruments, myRIO-1900) for a sampling time of 30 seconds. The experiments show the maximum velocity is located at the centerline; the profile decays with radial distance. The maximum velocity of the device is ~0.28 m/s for both devices/ fans and flowrate is ~6.3+/-0.7 slpm. Data in SI Figure S3.

**Bacteriophage MS2 propagation (Figure 8):** *Escherichia coli* bacteriophage MS2 was propagated using the Adam's Overlay method.[1] Briefly, Tryptic Soy Agar (TSA) plates were prepared and stored at RT overnight. A 2 mL aliquot of an overnight culture of the host species (*E. coli* Famp) was inoculated with 100 µL bacteriophage MS2 and incubated for 1 hr at 37 ºC with shaking. An additional 2-3 mL of overnight host culture was then mixed with the 1 hr phage MS2 culture and top agar before being poured onto prepared TSA plates for an overnight incubation at 37 ºC. Plates were then washed with 5-10 mL PBS for 5-10 mins before top agar was collected, rinsed with chloroform, and centrifuged (4000 x g for 15 mins at 4 ºC). Supernatant was then collected and mixed with glycerol for a final composition of 20% glycerol, 80% bacteriophage MS2 before being aliquoted. 1 mL aliquots were stored at -80 ºC until use.

(1) Adams, M.H. *Bacteriophages*. New York: Interscience Publishers, **1959**.

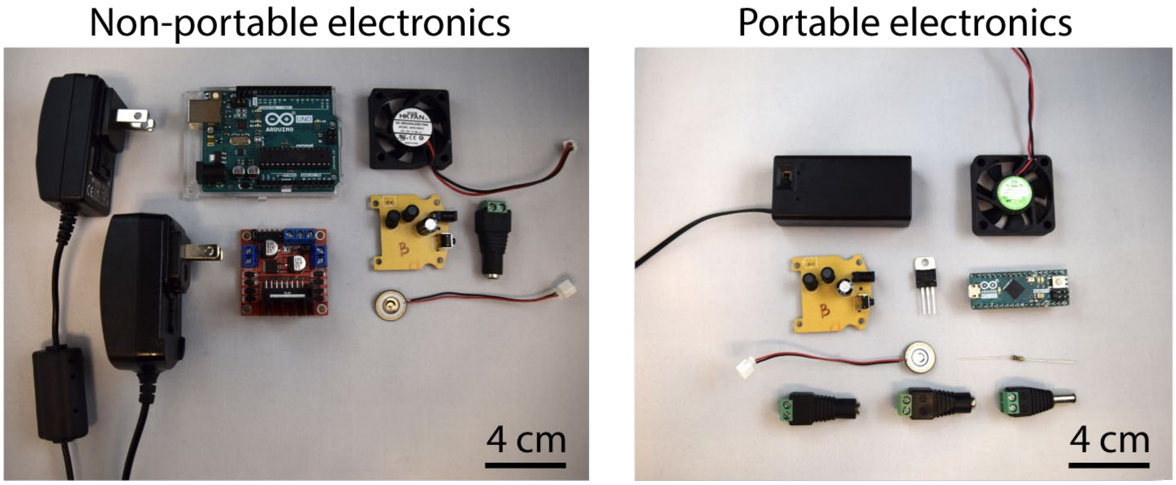


**Figure S1.** Photo of non-portable electronic components used for droplet retention measurements and videos (Figures 3-6) and portable electronics used for particle chamber experiments (Figure 7). Wires not pictured.

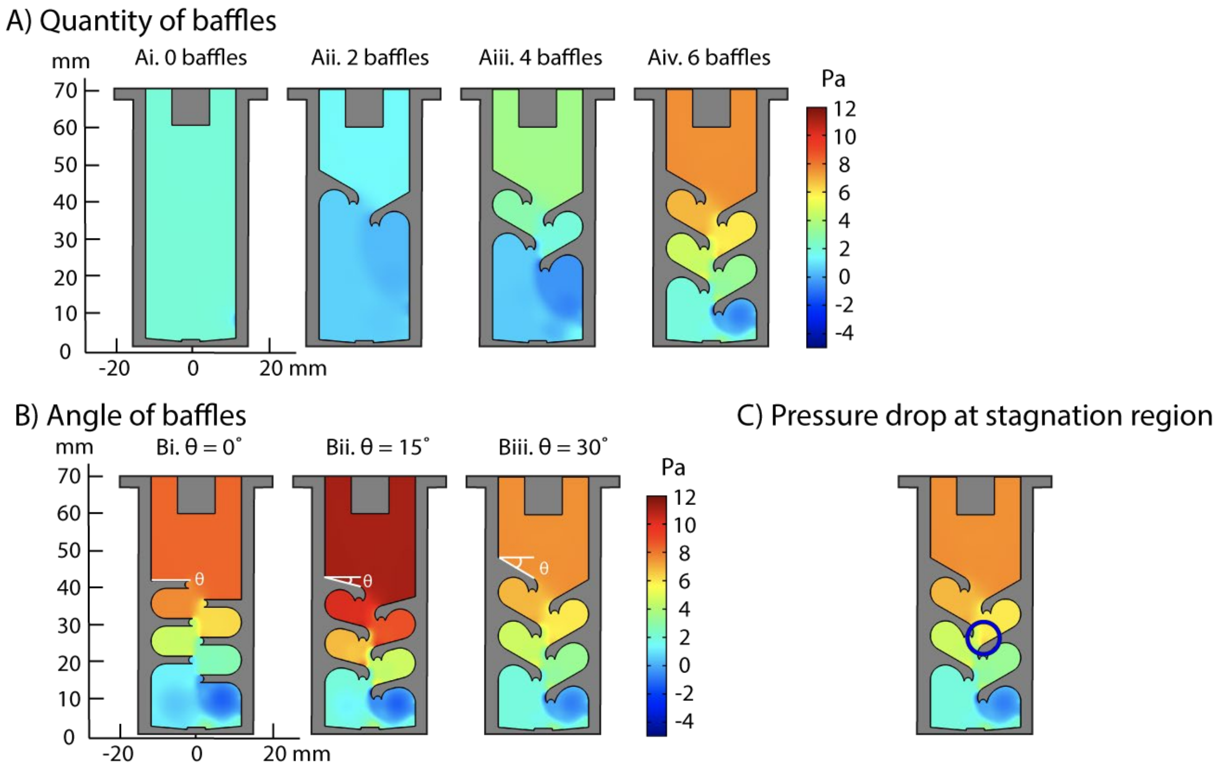


**Figure S2.** Pressure maps from simulations of device iterations pictured in Figure 5. (A) Pressure above the top baffles increases with each additional set of baffles. (B) When the angle of the baffle is changed from 0° to 15°, pressure increases before decreasing again at a 30° baffle angle. Reduced pressure is preferable to allow aerosol entrance into the device. (C) The pressure drop generated by impinging jets at stagnation regions increases droplet coalescence on baffle surfaces. Scale bar indicates the pressure above or below atmospheric pressure (i.e., 0 Pa = atmospheric pressure).

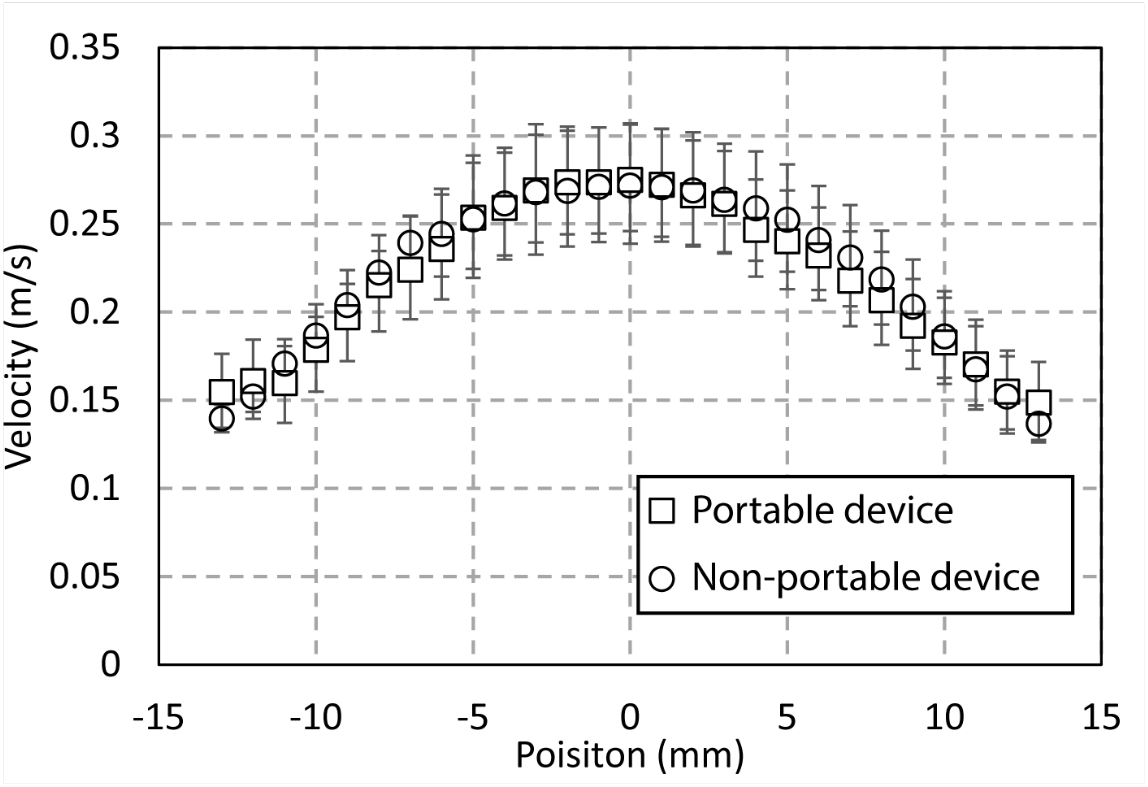


**Figure S3.** Plot of air flow velocity from the outlet of the device comparing the airflow rate between our non-portable device (Figures 3-6) and our portable device (Figure 7). The two devices used two different fans to accommodate voltage requirements for each electronics set up, but produced the same velocity and volumetric flow rate.

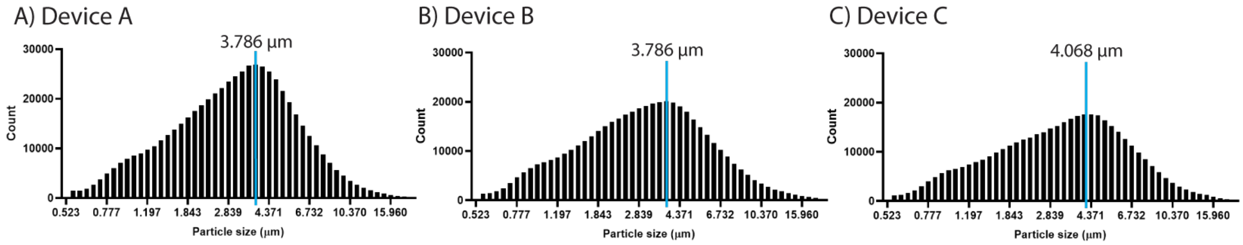


**Figure S4.** Droplet size distributions for each device as measured using an aerodynamic particle sizer. Average droplet size ± SD from n=3 devices is 3.88 ± 0.16 µm.

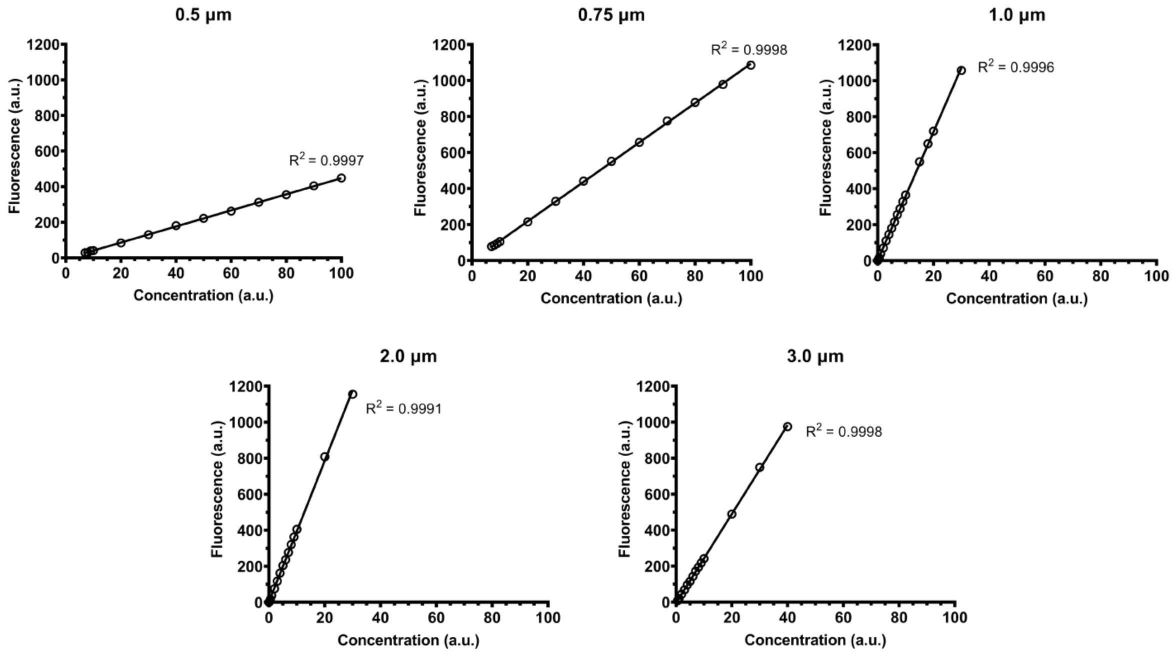


**Figure S5.** Calibration curves for each particle size used in Figure 7. Calibration curves were used to validate fluorescent signals of samples collected from devices and reference filters were in the linear range. Points represent mean ± SD of n=3. In all cases the standard deviation was smaller than the point plotted.

| Particle size (μm) | Concentration (particle/cm$^3$) |
|---|---|
| 0.50 | 4500-5500 |
| 0.75 | 10000-13000 |
| 1.0 | 4200-5800 |
| 2.0 | 1200-1500 |
| 3.0 | 1200-1500 |

**Table S1.** Particle concentration used in the test chamber for each particle size. Particle concentrations used at each size varied to ensure adequate particles were aerosolized and available for capture in the test chamber as in prior work (He, J.; Novosselov, I. V., Design and evaluation of an aerodynamic focusing micro-well aerosol collector. *Aerosol Science and Technology* **2017,** *51* (9), 1016-1026). During each 25-minute chamber run, 2-3 mL of solution was nebulized. Concentration reported here was measured with the aerodynamic particle sizer (APS) attached to the chamber exhaust.

| Diameter of inert particle proxy (μm) | Diameter of relevant bioaerosol (μm) | Reference | Examples of human health impact |
|---|---|---|---|
| 0.5 | SARS-CoV-2 aerosols<br>0.25 - 0.5 | Liu, Y. et al., *Nature* **2020**, *582* (7813), 557–560. | COVID-19, respiratory complications |
| 0.75 | *Staphylococcus aureus*<br>0.5 - 1.0 | Madsen, A. M. et al., *Annals of Work Exposures and Health* **2018**, *62* (8), 966–977. | Sepsis, pneumonia, infections |
| 1.0 | viral particle + respiratory droplet<br>≤ 1.0 | Fennelly, K. P. *The Lancet Respiratory Medicine* **2020**, *8* (9), 914–924. | Flu, common cold |
| 2.0 | *Aspergillus fumigatus*<br>2 - 3.5 | Kwon-Chung, K. J. et al., *PLoS Pathog* **2013**, *9* (12). | Pulmonary aspergillosis, allergic asthma |
| 3.0 | *Mycobacterium tuberculosis*<br>2 - 4 | Kim, K.-H. et al., *Journal of Environmental Sciences* **2018**, *67*, 23–35. | Tuberculosis |

**Table S2.** Diameter of aerosolized biological particles of interest. To avoid unnecessary exposure to known biological pathogens in this initial characterization publication, we chose to use monodispersed fluorescent polystyrene latex spheres as a proxy. Sizes used were based on relevant bioaerosols that are known to adversely affect human health.

A) 0.5 μm particles

| | **Device Avg (a.u.)** | **Device RSD (%)** | **Reference Avg (a.u.)** | **Reference RSD (%)** |
|---|---|---|---|---|
| **run 1** | 104.2 | **23.9** | 495.5 | **8.8** |
| **run 2** | 102.5 | **9.2** | 654.4 | **14.8** |
| **run 3** | 83.9 | **13.9** | 662.3 | **23.4** |

B) 0.75 μm particles

| | **Device Avg (a.u.)** | **Device RSD (%)** | **Reference Avg (a.u.)** | **Reference RSD (%)** |
|---|---|---|---|---|
| **run 1** | 164.7 | **25.8** | 1954.2 | **5.4** |
| **run 2** | 132.7 | **17.8** | 1138.3 | **19.0** |
| **run 3** | 107.0 | **19.8** | 1335.7 | **16.8** |

C) 1.0 μm particles

| | **Device Avg (a.u.)** | **Device RSD (%)** | **Reference Avg (a.u.)** | **Reference RSD (%)** |
|---|---|---|---|---|
| **run 1** | 68.3 | **56.6** | 1096.8 | **5.1** |
| **run 2** | 113.0 | **77.3** | 1206.0 | **14.8** |
| **run 3** | 66.7 | **17.0** | 880.2 | **19.4** |

D) 2.0 μm particles

| | **Device Avg (a.u.)** | **Device RSD (%)** | **Reference Avg (a.u.)** | **Reference RSD (%)** |
|---|---|---|---|---|
| **run 1** | 38.1 | **38.7** | 1237.1 | **7.6** |
| **run 2** | 55.3 | **16.2** | 1224.3 | **4.6** |
| **run 3** | 54.6 | **18.7** | 1366.9 | **4.9** |

E) 3.0 μm particles

| | **Device Avg (a.u.)** | **Device RSD (%)** | **Reference Avg (a.u.)** | **Reference RSD (%)** |
|---|---|---|---|---|
| **run 1** | 29.0 | **17.4** | 682.4 | **10.1** |
| **run 2** | 42.2 | **26.1** | 738.6 | **1.6** |
| **run 3** | 48.3 | **51.7** | 738.3 | **0.7** |

**Table S3.** Relative standard deviation of reference filters and devices across the three experimental test chamber runs. While variability is generally higher in our devices, it is important to note variability also exists within the reference filters used, likely due to the manual control of the flow rate and particle nebulization. The averages shown here were taken from n=3 devices or n=3 reference filters within a run of the test chamber. RSD=relative standard deviation.
